# GM1 and GD3 Gangliosides Attenuate NGF-TrkA and BDNF-TrkB Signaling Dysfunction Associated with Acute Diisopropylfluorophosphate Exposure in Mouse Brain

**DOI:** 10.1101/2025.03.31.646417

**Authors:** Yutaka Itokazu, Wayne D. Beck, Alvin V. Terry

## Abstract

The prevalence of neurodegenerative diseases and mental health disorders has been increasing over the past few decades. While genetic and lifestyle factors are important to the etiology of these illnesses, the pathogenic role of environmental factors, especially toxicants such as pesticides encountered over the life span, is receiving increased attention. As an environmental factor, organophosphates pose a constant threat to human health due to their widespread use as pesticides, their deployment by rogue militaries, and their use in terrorist attacks. The standard organophosphate-antidotal regimen provides modest efficacy against lethality, although morbidity remains high, and there is little evidence that it attenuates long-term neurobehavioral sequelae. Here we show that a novel intranasally administered treatment strategy with specific gangliosides can prevent the organophosphate-related alterations in important neurotrophin pathways that are involved in cognition and depression. We found that a single toxic dose of the organophosphate diisopropylfluorophosphate (DFP) in mice leads to persistent decreases in the neurotrophins NGF and BDNF and their receptors, TrkA and TrkB. Moreover, seven days of repeated intranasal administration of gangliosides GM1 or GD3 24 hours after the DFP injection prevented the neurotrophin receptor alterations. As NGF and BDNF signaling are involved in cognitive function and depression symptoms, respectively, intranasal administration of GM1 or GD3 can prevent the organophosphate-related alterations in those brain functions. Our study thus supports the potential of a novel therapeutic strategy for neurological deficits associated with a class of poisons that endangers millions of people worldwide.

**Highlights:** 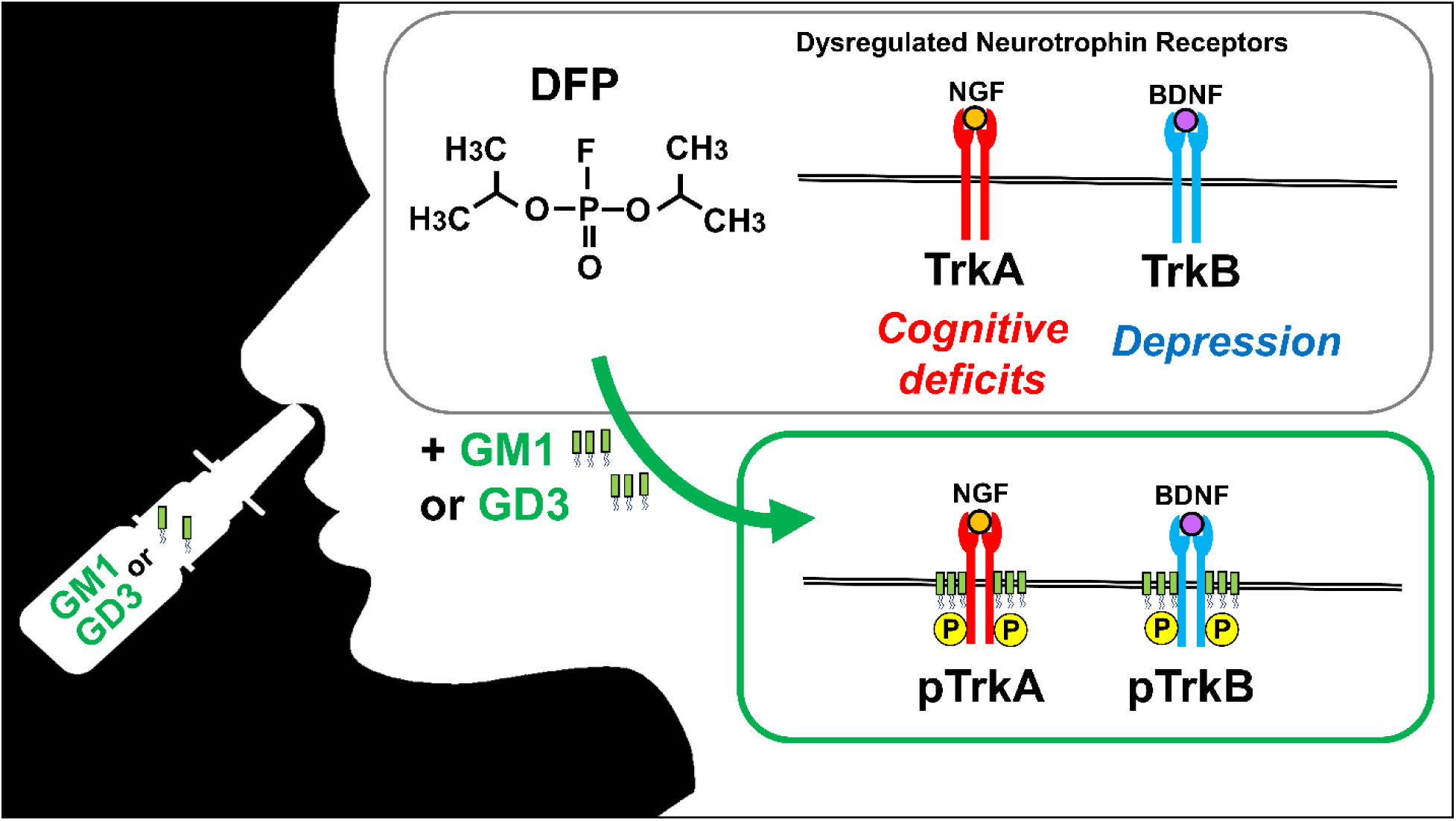

- A single exposure to DFP, which causes cognitive deficits, dysregulates NGF and BDNF signaling
- GM1 or GD3 24 hours after DFP injection prevents the alteration of the neurotrophin signaling
- Intranasal ganglioside treatment provides neuroprotective effects against persistent organophosphate toxicity

## 1. Introduction

The class of toxic chemicals known as organophosphates includes literally hundreds of compounds, most commonly used as pesticides. However, they are also found in a variety of other products, including defoliants, fire retardants, industrial solvents, lubricants, plasticizers, fuel additives, and chemical weapons/nerve agents (Costa 2018; Soltaninejad & Shadnia 2014). Regarding chemical weapons, the threat of intentional poisonings by rogue governments and terrorists, is an ever-present concern. Over the last 40 years, there have been multiple well-documented cases where organophosphate nerve agents were used against military soldiers and/or civilians. Examples include Iraqi military attacks on Iranian soldiers and Kurdish civilians in the 1980s, the Tokyo Sarin attacks by the domestic terrorist group Aum Shinrikyo in 1995, multiple military attacks on civilians in Syria with Sarin between 2013 and 2018, the use of the organophosphate VX in the assassination of the North Korean dictator’s half-brother in Malaysia in 2017, and the attempted assassinations of opponents of the Russian government with the organophosphate Novichok (e.g., the Skripals in 2018 in Britain and Alexei Navalny in Russia in 2020) (Naughton & Terry 2018; Steindl *et al*. 2021). In addition, there have been a number of accidental toxicant exposures during storage and shipments of hazardous chemicals over the years including a recent example, the Ohio train derailment in 2023. Acute, high dose organophosphate exposure from pesticides is also common in suicide attempts which continue to represent a major global health challenge (Huang *et al*. 2020; Mew *et al*. 2017).

The mechanism of the acute toxicity of organophosphates is the irreversible inhibition of acetylcholinesterase (AChE), which leads to marked elevations in synaptic acetylcholine levels. This, in turn, leads to excessive stimulation of cholinergic receptors (Pereira *et al*. 2014; Ecobichon 2001) which can result in seizures and death. The acute toxicity of organophosphates may also underlie (or contribute to) long-term neurological and neurobehavioral deficits in survivors which can include motor impairments, psychiatric disturbances, and cognitive impairments (Pereira *et al*. 2014). The standard organophosphate-antidotal regimen consists of respiratory support (mechanical ventilation), atropine (a muscarinic acetylcholine receptor antagonist), pralidoxime (an enzyme reactivator), and diazepam or midazolam (benzodiazepine anticonvulsants). This regimen provides modest efficacy against lethality, although morbidity remains high, and there is little evidence that it attenuates long-term neurobehavioral sequelae. Since both acutely toxic and repeated subacute exposures to organophosphates in animals have been shown to result in sustained increases in reactive oxygen species (ROS) and oxidative stress, mitochondrial dysfunction, lipid peroxidation, and neuroinflammation (Tsai & Lein 2021; Guignet & Lein 2019), treatments that target these factors (e.g., antioxidants, anti-inflammatory drugs) have been explored, although none of these approaches have resulted in new treatments to date.

In previous studies in rats, we found that repeated exposures to the representative nerve agent organophosphate, diisopropylfluorophosphate (DFP), at doses not associated with acute toxicity (1.0 mg/kg dose, alternate day administration over 30 days) led to deficits in learning and memory tasks, as well as alterations in neurotrophin-related proteins and cholinergic markers in brain regions important for cognition (Terry *et al*. 2011). Given their established roles in neuronal plasticity (i.e., both synaptic and morphological plasticity) (McAllister *et al*. 1999) neurotrophins and their receptors have been studied in a variety of neurodegenerative diseases and viewed as potential targets for drug discovery and development for several years. Of the various neurotrophins, nerve growth factor (NGF) may be especially important in age-related cognitive disorders given evidence of its decrease in the brain with age particularly in memory-related areas such as the hippocampus (Williams *et al*. 2006; Gomez-Pinilla *et al*. 1989; Larkfors *et al*. 1987). NGF is known to be especially important for the survival of forebrain cholinergic neurons (Counts & Mufson 2005) which are well documented to be involved in cognitive function, to degenerate with age, and to be markedly diminished in neurodegenerative illnesses such as Alzheimer’s disease (AD) brains (Bartus 2000). Additional support for the importance of NGF as a potential therapeutic target is evident in the results of experiments which suggested that deficits in NGF release and subsequent signaling (i.e., tyrosine receptor kinase phosphorylation) contribute to age-related deficits in long-term potentiation (Kelly *et al*. 2000), a form of neuronal plasticity that is widely believed to facilitate learning and memory (Bliss & Collingridge 1993). Under normal conditions, mature NGF binding to its high affinity receptor, Tropomyosin receptor kinase A (TrkA) promotes TrkA autophosphorylation which activates pathways that enhance cholinergic neuron survival (Counts & Mufson 2005). Conversely, proNGF, the uncleaved precursor form of NGF, binds to the tumor necrosis factor superfamily receptorp75 neurotrophin receptor (p75^NTR^) with higher affinity than mature NGF and it is more selective for the p75^NTR^ receptor relative to TrkA (Lee *et al*. 2001). Notably, the p75^NTR^ receptor is well-known for its role in mediating neuronal cell death (Underwood & Coulson 2008). There is also evidence that proNGF forms a heterotrimeric complex with the p75^NTR^ receptor and sortilin (a neurotensin receptor associated with intracellular sorting and trafficking of a variety of proteins) forming a high affinity binding site for proNGF to activate apoptotic cascades (Al-Shawi *et al*. 2008; Arnett *et al*. 2007; Nykjaer *et al*. 2004). This series of events may become more predominant in the setting of advanced age and neuropathological conditions such as AD. Al-Shawi and colleagues (Al-Shawi *et al*. 2008) concluded (based on extensive rodent experiments) that in old age, increases in the expression of sortilin enhance the vulnerability of neurons to age-related increases in pro-NGF, thus leading to neuronal loss and neurodegeneration. One objective of the experiments described here was, therefore, to evaluate the effects of DFP on the levels of NGF-related proteins in the rodent brain including proNGF, mature NGF, the neurotrophin receptors, TrkA, phospho-TrkA (i.e., the activated form of TrkA), p75^NTR^, and sortillin.

In this study, we were also interested in brain derived neurotrophic factor (BDNF), its precursor proBDNF, as well as the activated (i.e., phosphorylated) form of the BDNF receptor TrkB. BDNF signaling has been studied extensively in relation to several psychiatric disorders (including major depression) over the last several years due to its critical role in development, synaptic plasticity, cognitive function, and neuronal cell health (reviewed, Autry and Monteggia, 2012) (Autry & Monteggia 2012). Signaling by BDNF depends on the proteolytic cleavage of a pro-form of BDNF (proBDNF) to the mature form BDNF and these two proteins (like NGF and proNGF) can have opposing functions in the brain. For example, proBDNF binds preferentially to the p75^NTR^, mediating impairments in neuronal plasticity while facilitating hippocampal long-term depression, apoptosis and neuronal death (Barker 2009; Woo *et al*. 2005; Lee *et al*. 2001). Conversely, mature BDNF binds to TrkB and stimulates downstream signaling pathways leading to multiple neurotrophic effects: neuronal differentiation, neurite outgrowth, neuronal survival, strengthening of synapses, neurogenesis, long-term potentiation, and learning and memory (Mitchelmore & Gede 2014; Cunha *et al*. 2010; Barker 2009). Our observations that DFP exposure led to persistent decreases in the BDNF/proBDNF ratio (see results below) were interesting given the results of some animal and human studies of stress, aggression, and depression (Tognoli *et al*. 2010; Ilchibaeva *et al*. 2015; Zhao *et al*. 2017), which support the argument that the ratio of these proteins may be a more sensitive index of the biological changes in the brain that influence the behavioral outcomes than BDNF or proBDNF alterations alone.

We recently reported persistent effects of a single exposure to DFP 4.0 mg/kg on cognitive function, cellular senescence, and proinflammatory cytokines in the mouse brain (Terry *et al*. 2024). In this study we observed modest, but persistent DFP-related impairments in spatial learning and working memory. Histological and molecular experiments indicated that DFP was associated with persistent alterations in several senescence markers and proinflammatory cytokines in brain regions that are relevant to the performance of the memory-related tasks (e.g., hippocampus, prefrontal cortex).

Gangliosides (Fig. S1) are sialic acid-containing glycosphingolipids known to play essential roles in cell-cell recognition, adhesion, signal transduction, and cellular migration, and are crucial in all phases of neurogenesis (Itokazu & Yu 2023; Itokazu *et al*. 2023; Itokazu & Terry 2024). Acetylcholine is the neurotransmitter of the cholinergic system and plays an important role in memory, attention, motivation, novelty seeking, and arousal. Dysfunction of the cholinergic system is clinically significant in several neurodegenerative diseases, especially dementia (Terry & Buccafusco 2003). Gangliosides have been reported to interact with neurotrophin receptors and support neuroprotective phenomena as well as many other essential functions (Ledeen & Wu 2015; Fantini & Barrantes 2009; Furukawa *et al*. 2019; Ballough *et al*. 1998) that are important to cholinergic neurons. For example, cholinergic innervations of the rat cortex were lesioned and intracerebroventricular infusions of GM1 successfully attenuated the reductions in choline acetyltransferase (ChAT) levels. This GM1 effect was similar to that of NGF infusion, while a combination infusion of NGF and GM1 had a synergistic effect to significantly increase cholinergic presynaptic terminal size (Garofalo *et al*. 1993). In non-human primates, administration of NGF in combination with GM1 induced a long-term protective effect on nucleus basalis cholinergic neurons after neocortical infarction. (Liberini *et al*. 1993). The current data revealed significant decreases in TrkA and TrkB signaling in the cortex, prefrontal cortex, and hippocampus in DFP (4.0 mg/kg) exposed mice. Recently, we discovered administration of both GM1 and GD3 at doses up to 5.0 mg/kg are both safe (i.e., without adverse reactions) and effective in Parkinson’s disease model mice when administered by the intranasal route (Fuchigami *et al*. 2023; Itokazu *et al*. 2021).

Here, we evaluated a novel noninvasive intranasal treatment strategy against persistent organophosphate toxicity using specific gangliosides known to be play essential roles neuronal function and to have neuroprotective properties. We found that intranasal administration of gangliosides GM1 or GD3 24 hours after the DFP injection once daily for 7 days prevented the neurotrophin receptor alterations.

## 2. Materials and Methods

The time-line diagram of the experiment is displayed in Fig.1.

**Fig. 1.**
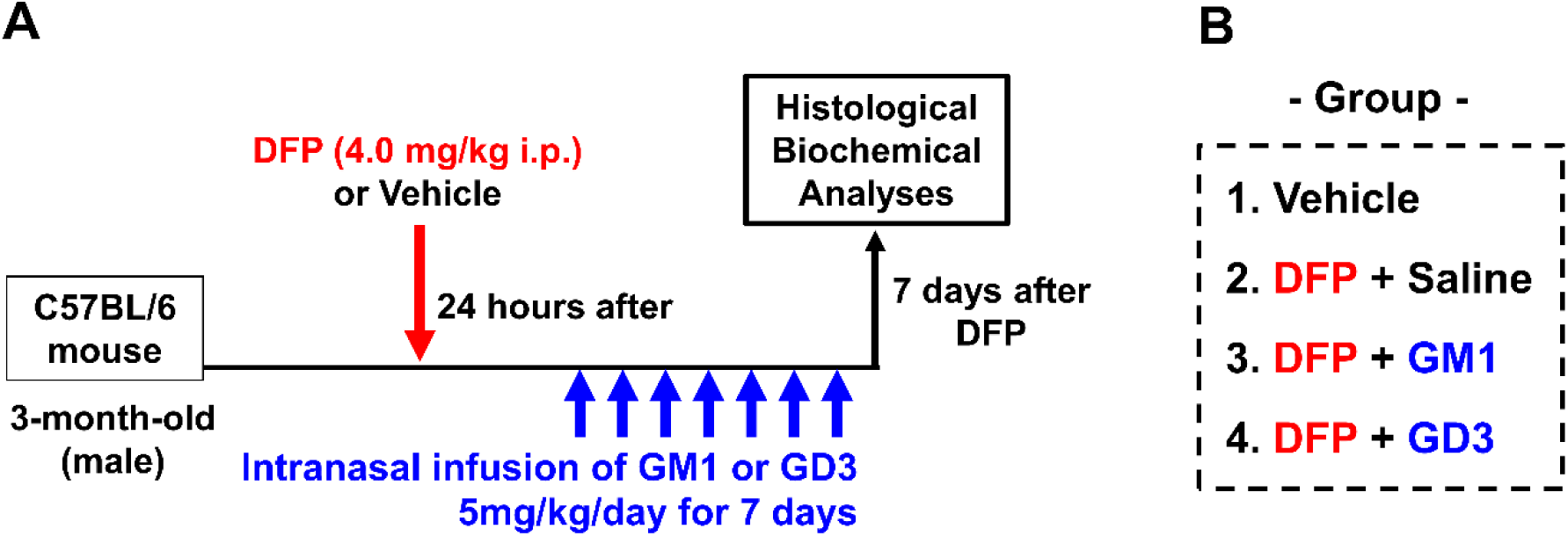
Time-line Diagram of the Experiment. (A) DFP at a dose of 4.0 mg/kg was injected (i.p.) into three-month-old mice. Gangliosides (GD3 or GM1; 5 mg/kg/day) were intranasally administered to DFP-exposed mice for 7 days, beginning 24 hours after the DFP injection. (B) Animals were divided into four groups: (1) vehicle group (n=6), (2) DFP-exposed mice with saline infusion as the placebo group (n=6), (3) DFP-exposed mice receiving GM1 (5 mg/kg/day) infusion (n=4), and (4) DFP-exposed mice receiving GD3 (5 mg/kg/day) infusion (n=4).

### 2.1. Antibodies

For this study, the following antibodies were purchased: rabbit anti-TrkA (RRID:AB_310180; Millipore Millipore, St. Louis, MO, USA, #06-574), mouse anti-NeuN (RRID:AB_2298772; Sigma-Aldrich, St. Louis, MO, USA, #MAB377), rabbit anti-p75 neurotrophin receptor (p75^NTR^, RRID:AB_310649; Millipore #07-476), rabbit anti-sortilin (RRID:AB_2192606; Abcam, Cambridge, MA, USA, #ab16640), goat anti-TrkB (RRID:AB_2155263; R&D Systems, Minneapolis, MN, USA, #BAF1494), rabbit anti-actin (RRID:AB_476693; Sigma #A2066), rabbit anti-NGF (RRID:AB_2040019, Alomone Labs, Jerusalem, Israel, #AN-240), rabbit anti-brain-derived neurotrophic factor (BDNF) (RRID:AB_2039756, Alomone Labs #ANT-010), and rabbit anti-phospho-Trk (RRID:AB_2298805; Cell Signaling Technology, Danvers, MA, USA, #9141).

### 2.2. Animals

All animal experiments were approved by the Institutional Animal Care and Use Committee (IACUC) at Augusta University (AU) according to the National Institutes of Health (NIH) guidelines and were performed with approved animal protocols (references AUP 2013-0526 and 2014-0694). Measures were taken to minimize pain or discomfort in accordance with the Guide for the Care and Use of Laboratory Animals, 11th edition, National Research Council, 2011. Significant efforts were also made to minimize the total number of animals used while maintaining statistically valid group sizes. Three-month-old, male C57BL/6 mice were obtained from Envigo, Indianapolis, IN, and housed at Augusta University in a temperature-controlled room (25°C), maintained on a 12-h light/dark cycle with free access to food (Teklad Rodent Diet, Inotiv, Madison, WI) and water throughout the study.

### 2.3. Diisopropylfluorophosphate administration

In previous studies in rats, we found that repeated exposures to DFP at doses not associated with acute toxicity led to deficits in learning and memory tasks, as well as alterations in neurotrophin-related proteins and cholinergic markers in brain regions important for cognition (Terry *et al*. 2011). In this report, we evaluated a 4.0 mg/kg dose of DFP for effects on neurotrophic signaling, as this dose was used in our most recent study (Terry *et al*. 2024), which resulted in persistent cognitive impairments. Each experimental mouse received vehicle (0.9% Blood Bank Saline, Thermo Fisher Scientific, Rockford IL, USA) or DFP, CAS 55–91-4 (Sigma Aldrich D0879-1G Lot: 126 K1306, St. Louis, MO) 4.0 mg/kg dissolved in vehicle (normal saline) and administered by intraperitoneal injection in a volume of 10x of body weight.

Individual mice were monitored (in their home cages for a period of approximately 5 minutes each day) for visible cholinergic signs (diarrhea, excessive salivation or lacrimation, respiratory difficulties, muscle fasciculations) or other signs of distress throughout the study.

### 2.4. Intranasal ganglioside administration

Currently, intracerebroventricular administration is the most reliable method for delivering gangliosides into the brain. We successfully developed a more convenient, non-invasive delivery procedure by intranasal infusion of gangliosides (Fuchigami *et al*. 2023; Itokazu *et al*. 2021). According to our previous experiments, we chose 5 mg/kg/day as the dose for intranasal ganglioside treatment without any anesthetic treatment. Gangliosides (GD3 or GM1; 5 mg/kg/day) were intranasally administered into DFP-exposed mice for 7 days beginning 24 hours after the DFP injection, using capillary tips (Bio-Rad, Hercules, CA, #2,239,915) with 6 μL administered into each of the right and left nares twice daily (6 μL × 2 nares = total 24 μL per day for 28 days). The GD3 and GM1 used in this study were isolated from either bovine buttermilk or brains in our laboratory by established procedures (Ariga *et al*. 1994; Itokazu *et al*. 2024; Ledeen & Yu 1982; Ren *et al*. 1992). GD3 and GM1, being amphipathic, were easily dissolved in saline. Animals were divided into four groups (Fig. 1): (1) vehicle group, (2) DFP-exposed mice with saline infusion as the placebo group, (3) DFP-exposed mice with GM1 (5 mg/kg/day) infusion, and (4) DFP-exposed mice with GD3 (5 mg/kg/day) infusion. Each group consisted of n = 4–6 animals.

### 2.5. Immunohistochemistry and imaging

Seven days after DFP or saline exposure, test subjects were anesthetized with isoflurane using an open-drop method and transcardially perfused with phosphate-buffered saline (PBS, pH 7.4) and 4% paraformaldehyde (PFA) in PBS. After perfusion, the brains were removed and post-fixed in 4% PFA at 4°C for 24 hours. The blocks were then equilibrated in sucrose (30% in PBS). Cryosections were cut at a thickness of 30 μm. Sections from the cortex were blocked and permeabilized with 1% Tween 20 in 1.0% bovine serum albumin (BSA) in PBS, at room temperature for 30 min, then washed with PBS before immunohistochemical staining. Sections were incubated in the following primary antibodies 37°C for 1 hour: rabbit anti-TrkA (1:100) and mouse anti-NeuN (1:250). After three washes with PBS, the samples were incubated with appropriate secondary antibodies coupled to Alexa488 or Alexa568 (Invitrogen) at a dilution of 1:1,000 at room temperature for 2 h. Nucleus counterstaining was performed with 1 μg/mL 40,6-diamidino-2-phenylindole (DAPI) (Thermo Fisher Scientific, #D1306) for 30 min. After washing with PBS three times, the sections were mounted with Vectashield mounting medium (Vector Laboratory, #H-1000). Confocal images were acquired using a Zeiss LSM 780 with a 40x oil objective (Zeiss, Land Baden-Württemberg, Germany) with identical acquisition settings.

### 2.6. Western blotting

The cortex, prefrontal cortex, hippocampus, and subventricular zone regions were isolated by dissection under a SZX7 stereo microscope (Olympus, Tokyo, Japan). Tissue blocks were lysed in radioimmunoprecipitation assay (RIPA) buffer containing 50 mM Tris-HCl, 150 mM NaCl, 5 mM NaF, 1 mM Na3VO4, 1% Nonidet P-40 (NP-40), 0.5% sodium deoxycholate, and 1% SDS (pH 7.5), supplemented with a complete protease inhibitor cocktail (Roche Applied Science, Indianapolis, IN, USA). Protein concentrations were measured using a bicinchoninic acid (BCA) protein assay kit (Thermo Fisher Scientific). Proteins were separated by SDS-PAGE (8% or 10% gel) under reducing conditions and transferred to polyvinylidene fluoride (PVDF) membranes. The membranes were probed with primary antibodies for TrkA (1:250), p75^NTR^, sortilin (1:3,000), TrkB (1:5,000), NGF (1:200), BDNF (1:400), Phospho-Trk (1:200), and actin (1:4,000), followed by appropriate secondary antibodies conjugated with horseradish peroxidase (anti-rabbit: BD Biosciences, San Jose, CA, USA, #554021; anti-goat: Santa Cruz Biotechnology, CA, USA, #sc-2020). Signals were visualized with Western Lightning western blot chemiluminescence reagent (PerkinElmer Life and Analytical Sciences, Waltham, MA, USA). The bands were quantified using the NIH ImageJ software (National Institutes of Health, Bethesda, MD, USA). Densitometric values were normalized by setting the proteins of interest, protein/actin, Trk/p75^NTR^, or phospho-Trk/Trk ratios for vehicle treatment in each condition. The normalized value from vehicle control was defined as 1.0.

### 2.7. Statistical analysis

To estimate biological variability, ensure the quality and consistency between biological samples, and maximize cost-effectiveness, we chose to use 4-6 animals per condition for ganglioside infusion experiments (1. vehicle group, n=6; 2. DFP-exposed mice with saline infusion as the placebo group, n=6; 3. DFP-exposed mice with GM1 infusion, n=4; and 4. DFP-exposed mice with GD3 infusion, n=4), based on our published study. All statistical procedures were performed using GraphPad Prism 10 (GraphPad, San Diego, CA, USA). The normality and homogeneity of variances of datasets were checked by a Kolmogorov-Smirnov test and Brown-Forsythe test, respectively. When datasets passed these tests, a one-way ANOVA with Tukey’s multiple comparison test was performed. In all cases, p-values are shown in the figure legends, and p < 0.05 was considered significant. All graphs depict mean ± S.E.M.

## 3. Results

### 3.1. Immunohistochemical analysis of TrkA expression in DFP-exposed mice

We administered DFP at a dose of 4.0 mg/kg to three-month-old mice as a single intraperitoneal (i.p.) injection. The 4.0 mg/kg dose of DFP was chosen based on the work of Locker *et al*. 2017 and O’Callaghan *et al*. 2015. While this sublethal dose has been associated with seizure activity and cholinergic signs in mice, it has also been associated with prolonged neuroinflammation in the brain, as indicated by elevated levels of inflammatory cytokines (Locker *et al*. 2017; O’Callaghan *et al*. 2015). Most recently, we demonstrated that this 4.0 mg/kg dose of DFP was linked to persistent alterations in several senescence markers and proinflammatory cytokines in brain regions relevant to memory-related tasks (e.g., hippocampus, prefrontal cortex) (Terry *et al*. 2024). Furthermore, a single exposure to this dose was associated with modest, but persistent impairments in acquisition and recall in a water maze spatial learning task. It also led to working memory deficits in a spontaneous alternation version of the Y-Maze task. In this study, immunohistochemical analysis revealed that the 4.0 mg/kg dose of DFP resulted in decreased TrkA expression in the cerebral cortex, as expected (Fig. 2). Based on this finding, we further examined the levels of NGF- and BDNF-related proteins.

**Fig. 2.**
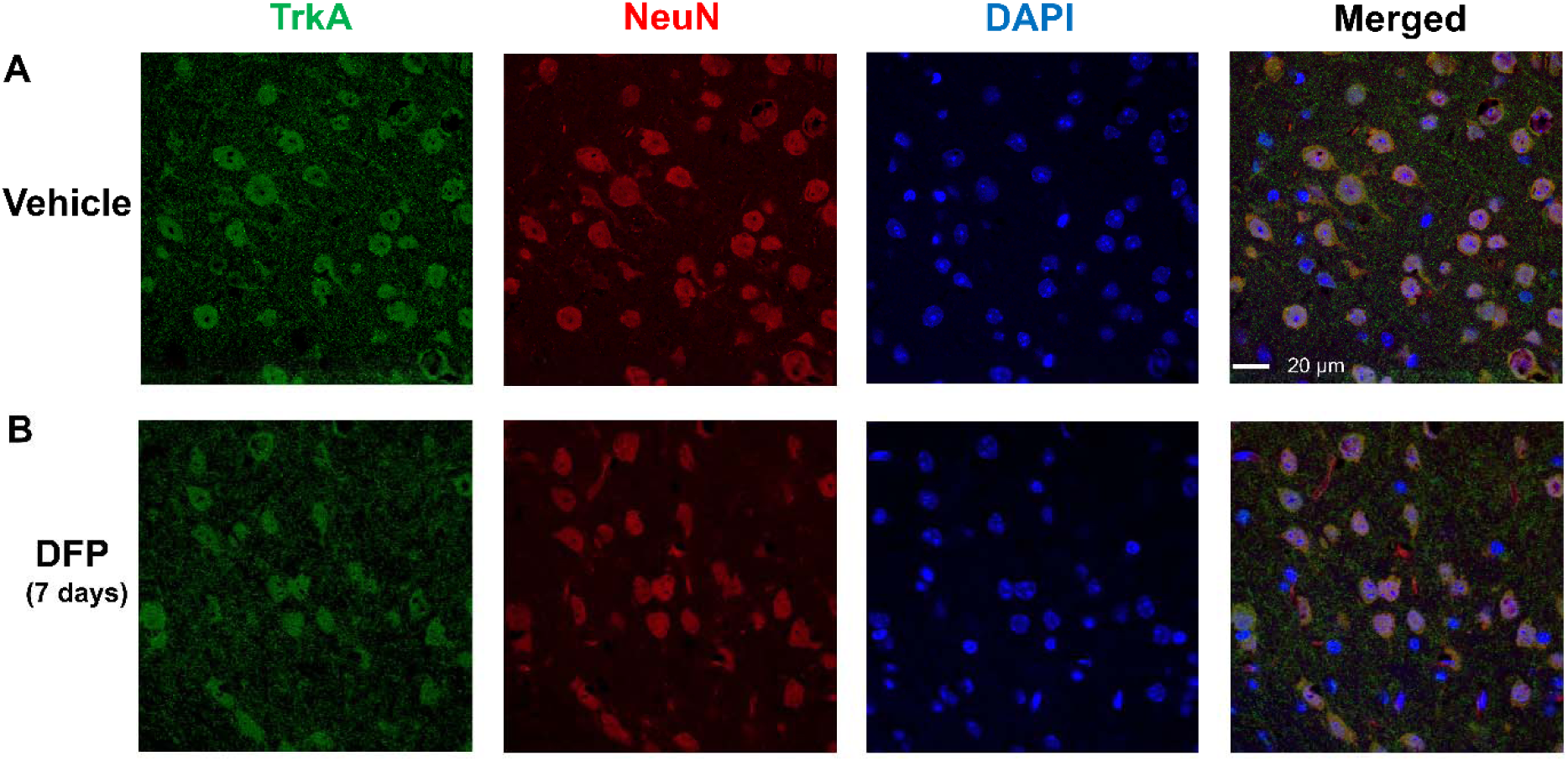
Altered TrkA Expression in the Cerebral Cortex Following DFP Exposure. The TrkA expression was examined in the cortex of mice 7 days after acute exposure to (A) vehicle or (B) DFP 4.0 mg/kg. Confocal immunohistochemistry images for TrkA (green), lineage-associated markers (red) for neurons (NeuN+ cells), nuclear DAPI staining (blue), and the merged images. Scale bars = 20 μm.

### 3.2. NGF- and BDNF-related proteins in DFP-exposed mice

Western blot analyses were performed using specific antibodies against TrkA, p75^NTR^, sortilin, and TrkB. Significant DFP-related decreases in TrkA expression were observed in the cortex, and in TrkB expression in the cortex and hippocampus (Fig. 3). On the other hand, significant increases in p75^NTR^ levels were detected in the prefrontal cortex and hippocampus after DFP exposure. It has been reported that GM1 binds to the Trk proteins and regulates receptor function (Mutoh *et al*. 1995). Therefore, we attempted intranasal ganglioside infusion to modulate NGF- and BDNF-related proteins, including TrkA and TrkB. Our previous studies demonstrated that intranasally administered GM1 was distributed to brain tissues, including the cortex, olfactory bulb, subventricular zone, hippocampus, midbrain, and cerebellum (Itokazu *et al*. 2023; Fuchigami *et al*. 2023; Itokazu *et al*. 2021). GM1 or GD3 was intranasally infused once per day for 7 days, beginning 24 hours after the DFP injection (Fig. 1). The intranasal infusion of gangliosides reversed the alterations in TrkA, p75^NTR^, and TrkB levels observed in the brains of DFP-injected mice (Fig. 3). The ratio of Trk to p75^NTR^ indicates whether a cell undergoes apoptosis or survives in the presence of neurotrophins (Terry *et al*. 2011). The ratio of TrkA/p75^NTR^ decreased in the cortex, prefrontal cortex, hippocampus, and subventricular zone in DFP-exposed subjects. Intranasal GD3 treatment restored the TrkA/p75^NTR^ ratio in the cortex, prefrontal cortex and hippocampus of DFP-affected mice. Intranasal GM1 treatment also restored these ratios in the cortex. Similarly, the ratio of TrkB/p75^NTR^ also decreased in the cortex, prefrontal cortex, hippocampus, and subventricular zone in DFP-exposed subjects. Intranasal GD3 treatment restored the TrkB/p75^NTR^ ratio in the prefrontal cortex and hippocampus of DFP-affected mice. Moreover, intranasal GM1 treatment also restored these ratios in the cortex and hippocampus.

**Fig. 3.**
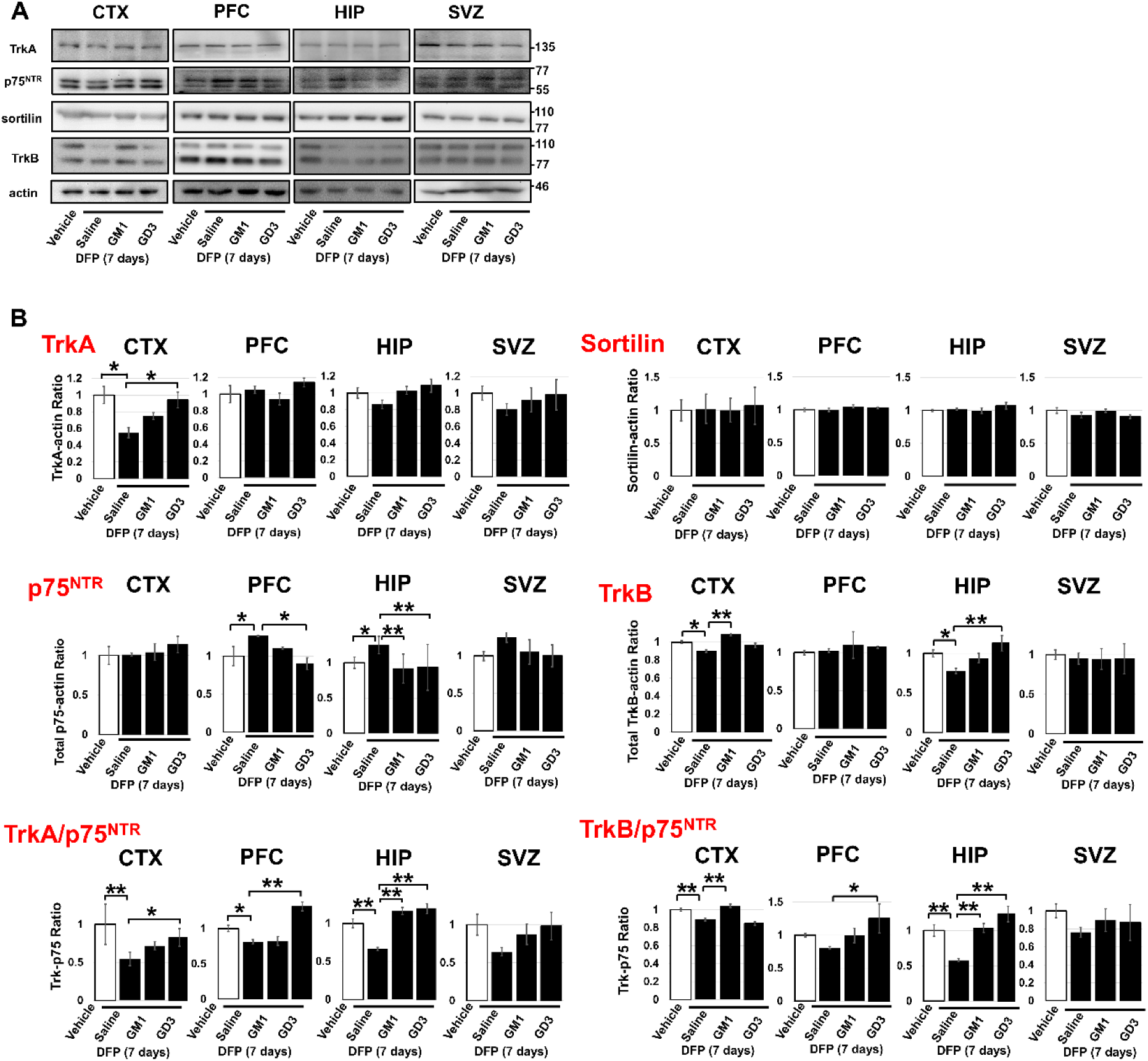
Expression of TrkA, p75^NTR^, Sortilin, and TrkB Following DFP Exposure and Ganglioside Treatment. TrkA and TrkB expression is reduced following DFP exposure (4.0 mg/kg) and restored by GM1 and GD3 treatments. Intranasally infused gangliosides (5 mg/kg/day for 7 days) ameliorate the decreased levels of TrkA and TrkB in the cortex, prefrontal cortex, and hippocampus of the DFP-exposed mouse brain. (A) Representative immunoblots. CTX, cortex; PFC, prefrontal cortex; HIP, hippocampus; SVZ, subventricular zone. (B) Signals on immunoblots were quantified through image analysis. Quantification of individual p75^NTR^ bands for each region can be found in Fig. S2. The columns represent the quantification of TrkA, TrkB, and Sortilin. Ratios of TrkA, TrkB, and Sortilin to actin were calculated for each replicate sample. The TrkA/p75^NTR^ and TrkB/p75^NTR^ ratios were analyzed to determine whether a cell is likely to undergo apoptosis or survives in the presence of neurotrophins. Quantification of individual bands of TrkB for each region can be found in Fig. S3. Results are means ± S.E.M. of ≥3 replicates per condition. *P < 0.05 and **P < 0.01 indicates significant differences between indicated groups, as calculated using one-way ANOVA with a Tukey’s multiple comparison test.

### 3.3. NGF and proNGF proteins

Western blot analysis using an antibody that recognizes both mature NGF and its precursor proNGF (14 kDa and 32 kDa immunoreactive bands, respectively, in Fig. 4A) indicated that NGF levels were decreased in the cortex, prefrontal cortex, and hippocampus of DFP-exposed mice compared to vehicle-treated mice (Fig. 4). In contrast, proNGF levels tended to increase in the cortex, prefrontal cortex, and hippocampus of DFP-exposed mice relative to vehicle-treated controls. The ratio of NGF to proNGF, which is critical for regulating cell survival and death, was significantly decreased in the cortex and prefrontal cortex, and showed a trend toward a decrease in the hippocampus of DFP-exposed mice compared to vehicle-treated mice. Treatment with GM1 restored the ratio of NGF to proNGF in the prefrontal cortex, and GD3 treatment restored it in the prefrontal cortex and hippocampus of DFP-injected mice.

**Fig. 4.**
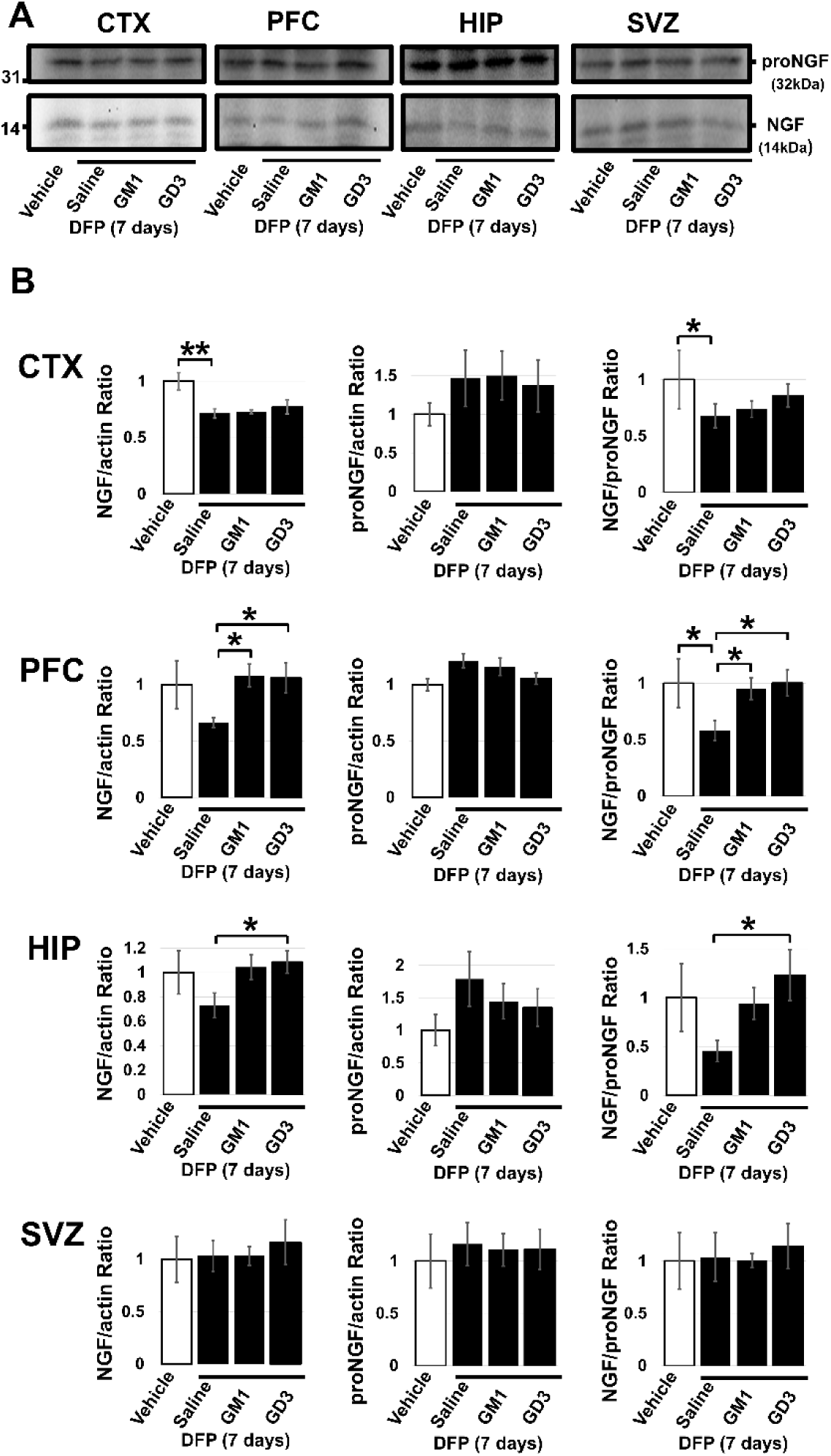
NGF and ProNGF Expression Following DFP Exposure and Ganglioside Treatment. Mature NGF and proNGF levels were measured by Western blot in brain lysates derived from mice exposed to either vehicle or DFP (4.0 mg/kg), followed by treatment with saline, GM1, or GD3 (5 mg/kg/day for 7 days). (A) Representative blot illustrating proNGF (32 kDa) and NGF (14 kDa). CTX, cortex; PFC, prefrontal cortex; HIP, hippocampus; SVZ, subventricular zone. (B) The data presented in the bar graphs were obtained from densitometry measurements of the bands for proNGF, NGF, and actin (Fig. 3). Ratios of proNGF and NGF to actin were calculated for each replicate sample. Results are expressed as means ± S.E.M. of ≥3 replicates per condition. *P < 0.05 and **P < 0.01 indicates significant differences between indicated groups, as calculated using one-way ANOVA with Tukey’s multiple comparison test.

### 3.4. BDNF and proBDNF proteins

We quantified precursor BDNF (proBDNF) and mature BDNF using Western blotting (14 kDa and 32 kDa immunoreactive bands, respectively, in Fig. 5A). The ratio of BDNF to proBDNF was significantly decreased in the cortex, prefrontal cortex, and hippocampus of DFP-exposed mice compared to vehicle-treated mice (Fig. 5). GM1 treatment restored the ratio of BDNF to proBDNF in the cortex and hippocampus of DFP-injected mice. GD3 treatment restored the levels of BDNF and proBDNF, as well as the ratio of BDNF to proBDNF in the hippocampus of DFP-affected mice.

**Fig. 5.**
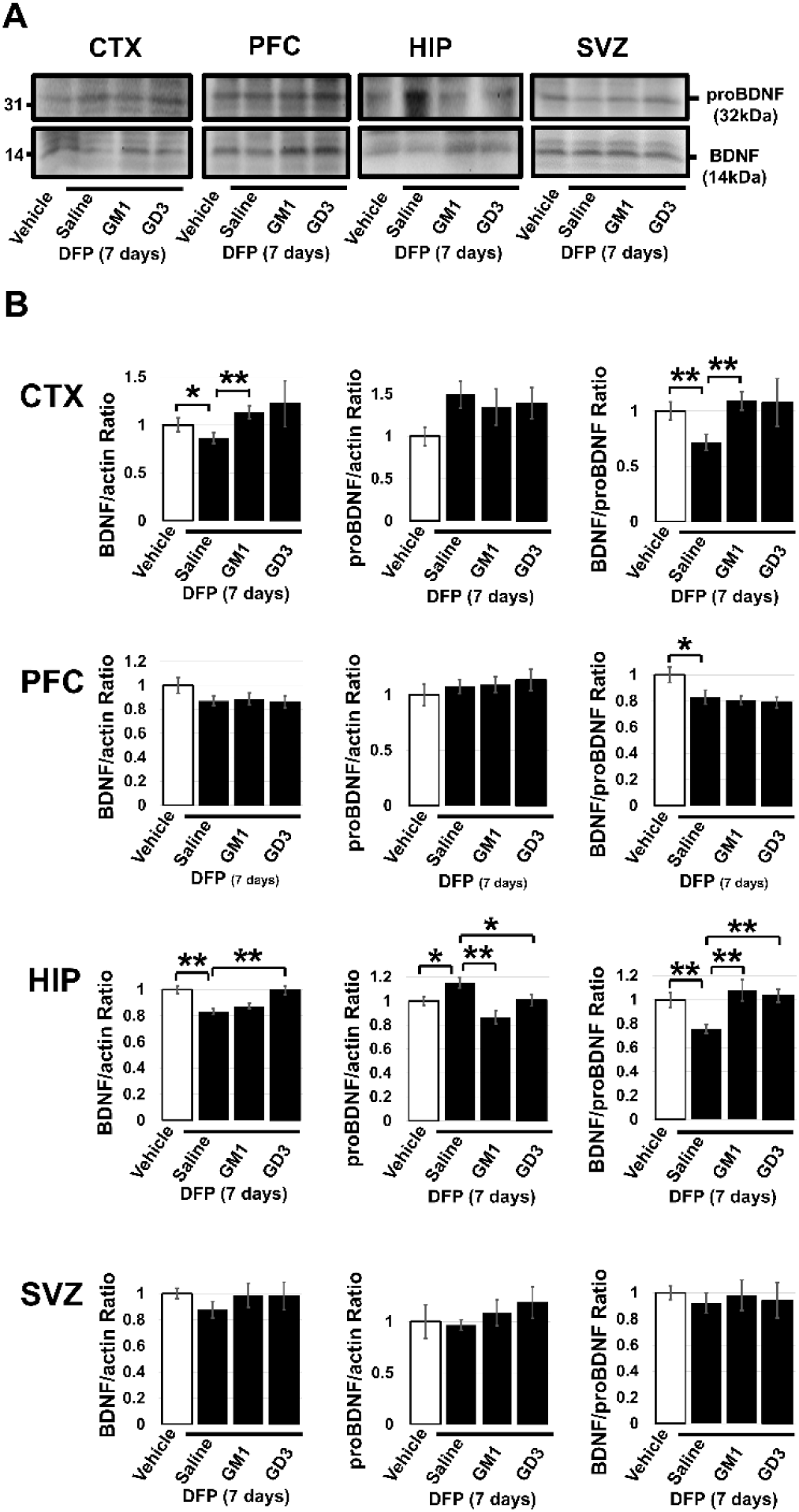
BDNF and proBDNF Expression Following DFP Exposure and Ganglioside Treatment. Mature BDNF and proBDNF levels were measured by Western blot in brain lysates derived from mice exposed to either vehicle or DFP (4.0 mg/kg), followed by treatment with saline, GM1, or GD3 (5 mg/kg/day for 7 days). (A) Representative blot illustrating proBDNF (32 kDa) and BDNF (14 kDa). CTX, cortex; PFC, prefrontal cortex; HIP, hippocampus; SVZ, subventricular zone. (B) The data presented in the bar graphs were obtained from densitometry measurements of the bands for proBDNF, BDNF, and actin (Fig. 3). Ratios of proBDNF and BDNF to actin were calculated for each replicate sample. Results are expressed as means ± S.E.M. of ≥3 replicates per condition. *P < 0.05 and **P < 0.01 indicates significant differences between indicated groups, as calculated using one-way ANOVA with Tukey’s multiple comparison test.

### 3.5. Phosphorylation of TrKA and TrkB

NGF signaling activation occurs through the binding of NGF to its receptor, TrkA, which induces phosphorylation. Similarly, TrkB activation is regulated by its autophosphorylation upon BDNF binding. The phosphorylated TrkA and TrkB receptors (pTrkA and pTrkB) were detected using a specific antibody. Levels of phosphorylated TrkA were significantly decreased in the cortex, prefrontal cortex, and hippocampus of DFP-exposed mice compared to vehicle-treated mice (Fig. 6). Likewise, phosphorylated TrkB levels showed significant decreases in the cortex and prefrontal cortex of DFP-exposed mice. Intranasal GM1 treatment restored the phosphorylation levels of TrkA in the cortex, prefrontal cortex, and hippocampus, and the phosphorylation levels of TrkB in the cortex and prefrontal cortex of DFP-injected mice. Intranasal GD3 treatment restored the phosphorylation levels of TrkA and TrkB in the cortex, prefrontal cortex, and hippocampus of DFP-injected mice. The phosphorylation of Trks and total Trk levels both changed after DFP exposure. Therefore, the values of phosphorylation were normalized by actin as an internal loading control (Fig. 6B and D). The data normalized by total Trks are shown in Fig. 6C and E.

**Fig. 6.**
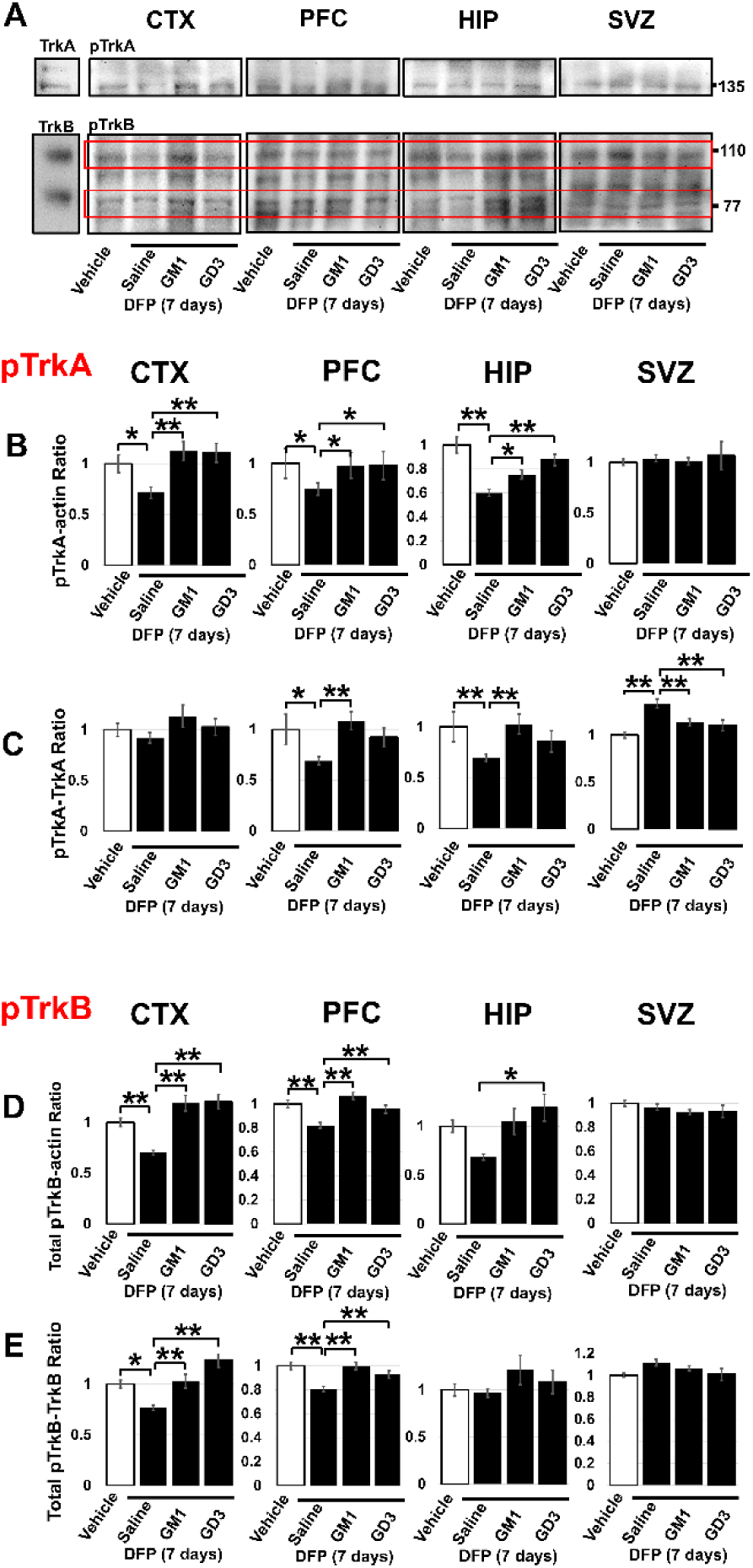
Phosphorylation of TrkA and TrkB Following DFP Exposure and Ganglioside Treatment. The phosphorylation of TrkA (pTrkA) and TrkB (pTrkB) is reduced following DFP exposure (4.0 mg/kg) and restored by GM1 and GD3 treatments. Intranasally infused gangliosides (5 mg/kg/day for 7 days) ameliorate decreased TrkA and TrkB levels in the cortex, prefrontal cortex, and hippocampus of the DFP-exposed mouse brain. (A) Representative immunoblots. The left bands exhibit total TrkA and TrkB bands from the lysate of the cortex from vehicle controls. CTX, cortex; PFC; prefrontal cortex, HIP, hippocampus; SVZ, subventricular zone. (B-D) Immunoblot signals were quantified through image analysis. Columns (B) and (D) represent pTrkA and pTrkB ratios to actin for each replicate. (C) pTrkA/TrkA and (E) pTrkB/TrkB ratios were analyzed. Quantification of individual pTrkB bands for each region can be found in Fig. S4. Results are means ± S.E.M. of ≥3 replicates per condition. *P < 0.05 and **P < 0.01 indicates significant differences among groups, calculated with one-way ANOVA and Tukey’s multiple comparison test.

## 4. Discussion

The major findings of this rodent study can be summarized as follows: (1) a single exposure to a 4.0 mg/kg dose of DFP in mice was associated with dysregulation of TrkA-NGF and TrkB-BDNF signaling, which are linked to cognitive function and depression; (2) administration of GM1 or GD3, 24 hours after the DFP injection and given once daily for 7 days, prevented alterations in neurotrophin signaling; and (3) this noninvasive intranasal treatment strategy using specific gangliosides shows promise as a novel approach for providing neuroprotective properties against persistent organophosphate toxicity.

In our recent report, we demonstated that this particular organophosphate dosing model (acute, relatively high dose of 4.0 mg/kg DFP exposure) led to persistent cognitive-domain-specific impairments, including deficits in spatial learning and recall as well as working memory (Terry *et al*. 2024). DFP exposure was also not associated with negative effects on weight or impairments in noncognitive (e.g., motor function or exploratory activity) behavioral assessments. Furthermore, we showed that DFP exposure was associated with persistent alterations in several senescence markers and proinflammatory cytokines in brain regions relevant to memory-related task performance (e.g., prefrontal cortex and hippocampus). Thus, single acute exposure to organophosphates like DFP leads to persistent impairments in specific domains of cognition that may be related to alterations in cellular senescence and inflammaging in the brain. In the current study, we specifically focused on the effects of these single exposures to DFP on neurotrophin signaling. Furthermore, we determined whether intranasal GM1 or GD3 could prevent persistent DFP-related alterations in the NGF and BDNF neurotrophin pathways.

DFP was associated with protracted (brain region-dependent) alterations in NGF-related proteins in brain regions known to support information processing and cognitive function (e.g. cortex, prefrontal cortex, hippocampus), indicating a potential mechanism for the behavioral effects we published recently (Terry *et al*. 2024). In previous studies in rats, we found that repeated exposure to DFP at doses not associated with acute toxicity led to deficits in learning and memory tasks as well as alterations in NGF-related proteins and cholinergic markers in brain regions important for cognition (Terry *et al*. 2011). The TrkA protein, a neurotrophin receptor normally associated with neuronal survival and plasticity was diminished in the cortex. We detected DFP-related elevations in neurotrophin proteins that have been associated with apoptosis and neuronal death (i.e. proNGF and p75^NTR^) in the prefrontal cortex and hippocampus. These observations are notable considering the deficits in performance of hippocampus-dependent memory tasks described recently. Given that the literature commonly describes proNGF-p75^NTR^ and NGF-TrkA as having opposing roles (apoptotic versus neurotrophic, respectively), we analyzed the TrkA/p75^NTR^ and NGF/proNGF ratios to predict whether cells in the brain are likely to undergo apoptosis or survival in the presence of neurotrophins. The ratios of TrkA/p75^NTR^ and NGF/proNGF decreased in the cortex, prefrontal cortex, and hippocampus in DFP-exposed subjects (Fig. 3). Intranasal ganglioside treatments restored the TrkA/p75^NTR^ and NGF/proNGF ratios in DFP-affected mice. It has been reported that a 0.5 mg/kg DFP injection for for 5 days resulted in decreased BDNF expression in the hippocampus, leading to a depression-like phenotype in the rats (Ribeiro *et al*. 2020). Our current western blot analysis also detected significant downregulation of BDNF levels in the cortex as well as hippocampus of DFP-exposed mice compared to vehicle control animals (Fig. 5). A potential mechanism for the reduced BDNF levels was suggested to involve epigenetic regulation, with decreased histrone acetylation at the BDNF promoter in the hippocampus of DFP-affected mice (Ribeiro *et al*. 2021). Both NGF and BDNF signaling are altered by DFP exposure, which may result in cognitive impairment and depression.

According to our studies and those of others, gangliosides have been considered neuroprotective molecules. GM1 binds to the Trk proteins and regulates their receptor function. GM1 specifically increases NGF-induced autophosphorylation of TrkA but not of p75^NTR^ (Mutoh *et al*. 1995). GM1 was able to prevent apoptotic cell death by GM1-activated Trk autophosphorylation (Ferrari *et al*. 1995). Trk receptors (TrkA, TrkB, TrkC) can be activated by GM1 in brain slices from the frontal cortex, hippocampus, and striatum (Duchemin *et al*. 2002). Exogenous GM1 (i.p.) increased BDNF levels in rat brains (Valdomero *et al*. 2015). Overexpression of GD3 synthase enhanced continuous phosphorylation and sustained dimerization of TrkA in the absence of NGF treatment in the cells (Fukumoto *et al*. 2000). In primary cerebellar granule cells, treatment with an anti-GD3 antibody inhibited BDNF-induced migration and attenuated BDNF-induced Akt activation, while the anti-GD1b antibody had no such effect (Komatsuya *et al*. 2022). GM1 and GD3 activate Trk receptors by enhancing phosphorylation and dimerization, thereby sustaining NGF and BDNF signaling

We previously reviewed literature suggesting that the administration of the GM1 may serve as a protective agent against organophosphate toxicity in rodents. (Itokazu & Terry 2024). It was previously shown that pretreatment with intracerebroventricular infusion of GM1 for 4 days reduced soman-induced seizure-related brain damage, while i.p. injection of GM1 failed to provide protection (Ballough *et al*. 1998). The blood–brain barrier (BBB) limits transportation of central nervous system (CNS) therapeutics. In previous studies, intracerebroventricular administration was the most reliable method of delivering gangliosides into the brain (Svennerholm *et al*. 2002; Itokazu *et al*. 2019; Galleguillos *et al*. 2022). We have successfully developed a more convenient noninvasive delivery procedure via intranasal infusion of gangliosides. Our most recent work indicates that both GM1 and GD3 at doses up to 5.0 mg/kg are both safe (i.e., without adverse reactions) and effective in Parkinson’s disease models when administered via the intranasal route (Fuchigami *et al*. 2023; Itokazu *et al*. 2021). In the work described in this manuscript significant DFP-related reductions in TrkA and TrkB in the cortex, prefrontal cortex, and hippocampus were observed. Furthermore, these DFP effects were prevented by intranasal ganglioside (GM1 and/or GD3) treatment administered 24 hours after the DFP injection, once daily for 7 days. Since it has been reported that less than 0.4% of intravenously injected GM1 can enter the non-human primate (monkey) brain (Revunov *et al*. 2020), the intranasal route is expected to result in significantly more ganglioside penetration into the CNS. Additionally, our intranasal delivery approach is much less invasive than other methods of delivering gangliosides to the brain (e.g., intracerebroventricular administration). Our novel, noninvasive intranasal treatment is a promising strategy for addressing persistent neurotrophic and neurobehavioral deficits associated with acute DFP exposure by leveraging the properties of specific gangliosides.

There is significant published literature that supports the link between ganglioside function and neuotrophic activity that is relevant to our findings in the this report. For example, the deletion of GD3 led to impairments in hippocampus-dependent memory functions and depression-like behaviors in mice (Fuchigami *et al*. 2024; Tang *et al*. 2021; Itokazu *et al*. 2018; Wang *et al*. 2014). GM1-knockout mice exhibit impaired movement and virtually all the neuropathological findings of Parkinson’s disease, including memory defects (Wu *et al*. 2020). DFP exposure resulted in cognitive impairments and depression symptoms (Ribeiro *et al*. 2020; Terry *et al*. 2024), similar to those observed in ganglioside knockout mice, but not motor function deficits. Our investigations into the physiological roles of GD3 and GM1 in postnatal neurogenesis revealed that GD3 augments neural stem cells, while GM1 enhances neuronal differentiation in mouse brains (Fuchigami *et al*. 2023; Itokazu *et al*. 2019). Notably, intracerebroventricular administration of GM1 to patients with AD was reported to halt the progressive deterioration of nerve processes and increase the turnover of neurotansmitters (Svennerholm *et al*. 2002). Regarding nuclear gangliosides, we discovered that GM1 promotes neuronal differentiation through an epigenetic regulatory mechanism. GM1 binds with acetylated histones on the promoters of GM2 synthase (GM2S; N-acetylgalactosaminyltransferase [GalNAcT]), a critical enzyme for GM1 synthesis, as well as on neuronal genes. Exogenous GM1 induces the transcription of GM2S by increasing histone acetylation levels at the GM2S promoter region, thereby recruiting more transcription factors. Consequently, more endogenous GM1 is synthesized following GM1 administration (Itokazu *et al*. 2021; Itokazu *et al*. 2016; Tsai *et al*. 2016; Tsai & Yu 2014). These ganglioside functions are valuable for developing novel strategies to treat cognitive and mental health diseases induced by organophosphates.

GM1 is known to bind to TrkA, enhancing TrkA’s response to NGF stimulation. This interaction results in higher autophosphorylation and increased dimerization (Farooqui *et al*. 1997; Mutoh *et al*. 1995). It has been reported that overexpression of GD3 synthase enhances continuous phosphorylation and sustains dimerization of TrkA (Fukumoto *et al*. 2000). Although the specific roles of GM1 and GD3 in protecting against DFP-induced dysregulation of neurotrophin signaling remain unclear, restoring Trk phosphorylation, GM1 seems more effective in the cortex, while GD3 has a greater impact in the hippocampus. In the prefrontal cortex, the selectivity of both gangliosides is intermediate between those observed in the cortex and hippocampus. GM1 is a major ganglioside in neurons, whereas GD3 is expressed in neural stem cells and immature neurons (Itokazu *et al*. 2018). The ratio of the number of neurons to non-neuronal cells is higher in the cortex than in the hippocampus, with the ratio in the frontal cortex being intermediate between that of the cortex and the hippocampus (Keller *et al*. 2018). It suggests that the cortex hosts more neurons affected by GM1, while the hippocampus contains more GD3-responsive cells, including neural stem cells, progenitor cells, and immature neurons.

Our previous research (Terry *et al*. 2024) demonstrated that the senescence marker p21 (also known as cdkn1a) showed a marked elevation in the prefrontal cortex after exposure to DFP (single dose of 4.0 mg/kg DFP). In a Parkinson’s disease model in mice, intranasal infusion of GD3 reversed the elevated p21 expression (Fuchigami *et al*. 2023). To determine whether ganglioside treatment (intranasal GM1 or GD3 for 7 days) might also restore p21 expression in DFP-exposed mice, we performed an initial, preliminary immunohistochemistry experiment. We found that intranasal GD3 treatment downregulated p21 levels, while GM1 treatment partially reduced these levels in the prefrontal cortex of DFP-exposed mice (Fig. S5). Future studies will be conducted to build on these preliminary findings. Interestingly, the alterations in p21 expression were observed in glial cells rather than neuronal cells (Terry *et al*. 2024). Astrocytes and microglia are known to express several neurotransmitter receptors, including α7 nicotinic acetylcholine receptors. Cholinergic pathways in glial cells may play an important role in contributing to the anti-inflammatory pathway in DFP-exposed mice. It has been reported that GM1 decreases inflammatory microglia and that both GM1 and GD3 increase microglial phagocytosis (Park *et al*. 2009; Schneider *et al*. 2022; Galleguillos *et al*. 2022; Wang *et al*. 2021). These are additional important roles of gangliosides: they modulate microglial inflammatory responses and facilitate the internalization of toxic proteins by microglia. Persistent inflammatory conditions are considered to contribute to neurodegenerative disorders. To address organophosphate-induced neurological deficits, targeting inflammation is a promising approach for the development of future therapies that may involve ganglioside administration.

Further experiments are in progress to elucidate the detailed mechanisms underlying the negative effects DFP in the brain and restorative effects of ganglioside discussed in this report. Nevertheless, it is safe to conclude from the results of this study that intranasal gangliosides improve NGF-TrkA and BDNF-TrkB signaling pathways in the cortex, prefrontal cortex, and hippocampus after DFP exposure. Our observations suggest that intranasal gangliosides GM1 or GD3 can prevent organophosphate-related alterations in important neurotropic pathways in the brain, which may in turn improve the cognitive impairments and depression-related symptoms asssociated with organophosphate exposure.

## CRediT authorship contribution statement

**Yutaka Itokazu**: Writing – original draft, Writing – review & editing, Methodology, Validation, Visualization, Investigation, Formal analysis, Conceptualization, Funding acquisition, Resources. **Wayne D. Beck**: Writing – review & editing, Methodology, Validation, Investigation. **Alvin V. Terry Jr.**: Writing – review & editing, Methodology, Validation, Conceptualization, Funding acquisition, Supervision, Project administration, Resources.

## Acknowledgments

We thank Drs. Dongpei Li and Toshio Ariga, and Truc Bui for excellent technical support, and Ashley Davis for her administrative assistance in preparing this manuscript. We would also like to acknowledge the Augusta University Cell Imaging Core facility (Director Graydon Gonsalvez, Core Manager Rachel Cui) for assistance with confocal microscope imaging experiments. The authors also wish to acknowledge the outstanding infrastructural support provided by the Department of Pharmacology & Toxicology, Medical College of Georgia at Augusta University.

## Data availability

Data will be made available on request.

## Supplementary Figures

**Fig. S1.**
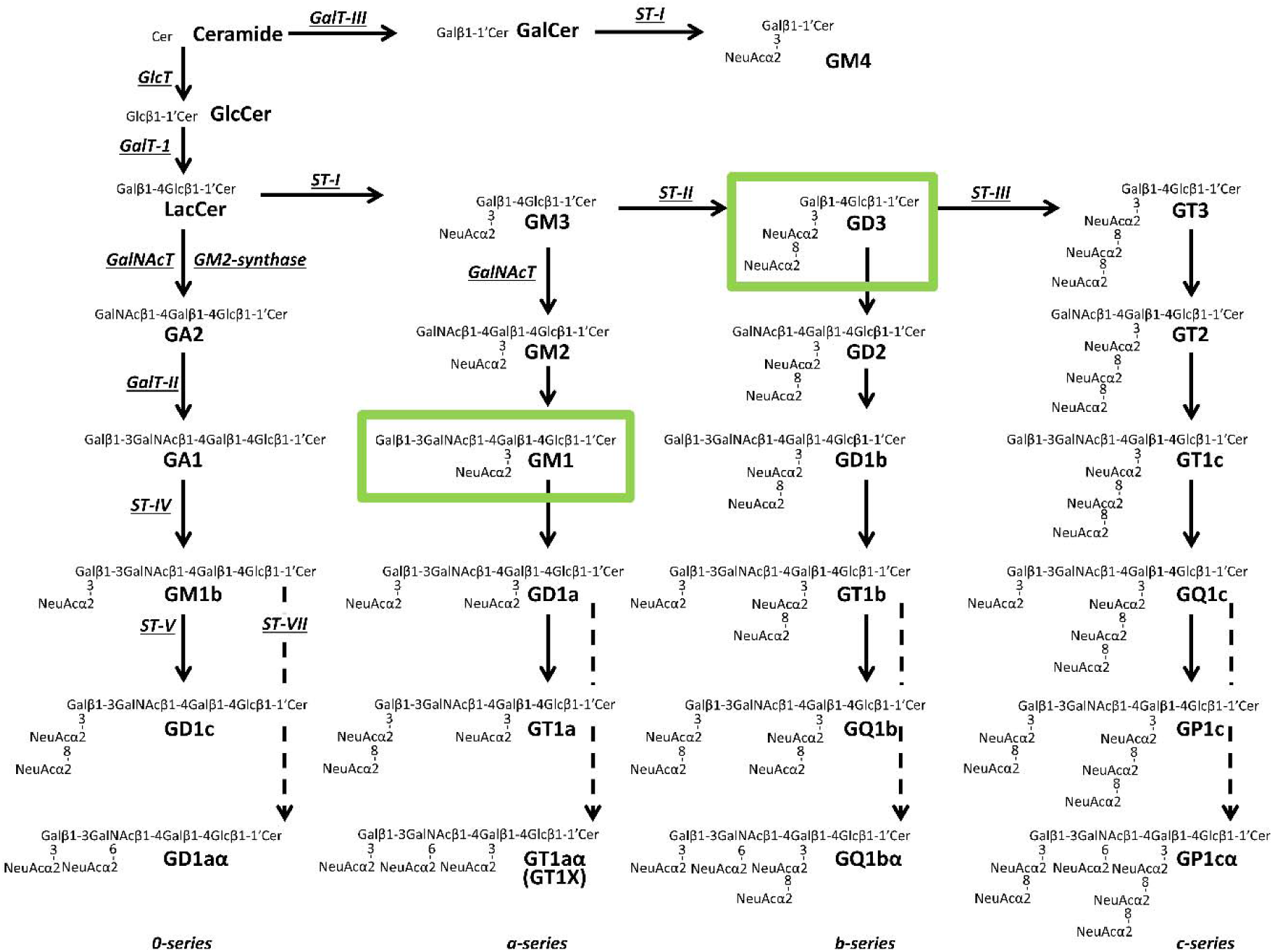
Metabolic pathways and structure of glycosphingolipids, including gangliosides. Cer, ceramide; GalNAc-T, *N*-acetylgalactosaminyltransferase I (***B4galnt1***, GA2/GM2/GD2/GT2-synthase); GalT-I, galactosyltransferase I (***B4galt6***, lactosylceramide synthase); GalT-II, galactosyltransferase II (***B3galt4***, GA1/GM1/GD1b/GT1c-synthase); GalT-III, galactosyltransferase III (***Ugt8a***, galactosylceramide synthase); GlcT, glucosyltransferase (***Ugcg***, glucosylceramide synthase); ST-I, sialyltransferase I (***St3gal5***, GM3/GM4-synthase); ST-II, sialyltransferase II (***St8Sia1***, GD3-synthase); ST-III, sialyltransferase III (***St8Sia3***, GT3-synthase); ST-IV, sialyltransferase IV (***St3gal2***, GM1b/GD1a/GT1b/GQ1c-synthase); ST-V, sialyltransferase V (***St8sia5***, GD1c/GT1a/GQ1b/GP1c-synthase); ST-VII, sialyltransferase VII (***St6galnac6***, GD1aα/GT1aα/GQ1bα/GP1cα-synthase). Official symbols of genes are represented in ***italics*** in this figure legend. GM1 and GD3 used in this study are marked in green triangles. GM1, GD1a, GD1b and GT1b are the most abundant ganglioside species in adult mammalian brain and neurons.

**Fig. S2.**
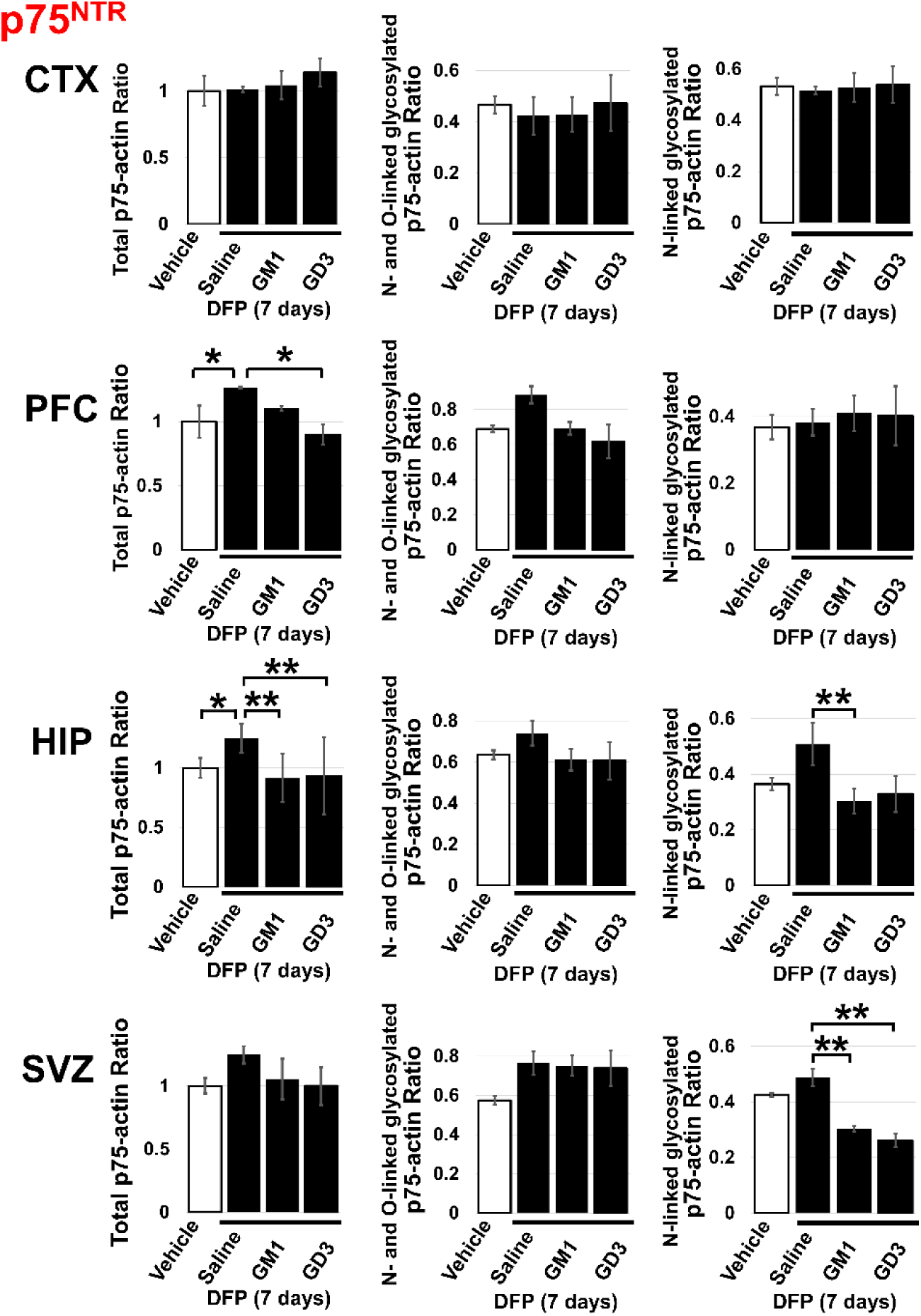
Quantification of Differentially Glycosylated p75^NTR^. p75^NTR^ expression are upregulated following DFP exposure (4.0 mg/kg) and restored by GM1 and GD3 treatments. Intranasally infused gangliosides (5 mg/kg/day for 7 days) ameliorate the decreased levels of p75^NTR^ in the prefrontal cortex, and hippocampus of the DFP-exposed mouse brain. Signals on immunoblots of Fig. 3A were quantified through image analysis. Quantification of upper (N-and O-linked glycosylation) and lower (N-linked glycosylation) p75^NTR^ are shown in the columns. The columns represent the quantification of p75^NTR^ to actin were calculated for each replicate sample. Results are means ± S.E.M. of ≥3 replicates per condition. *P < 0.05 and **P < 0.01 indicates significant differences between indicated groups, as calculated using one-way ANOVA with a Tukey’s multiple comparison test. CTX, cortex; PFC, prefrontal cortex; HIP, hippocampus; SVZ, subventricular zone.

**Fig. S3.**
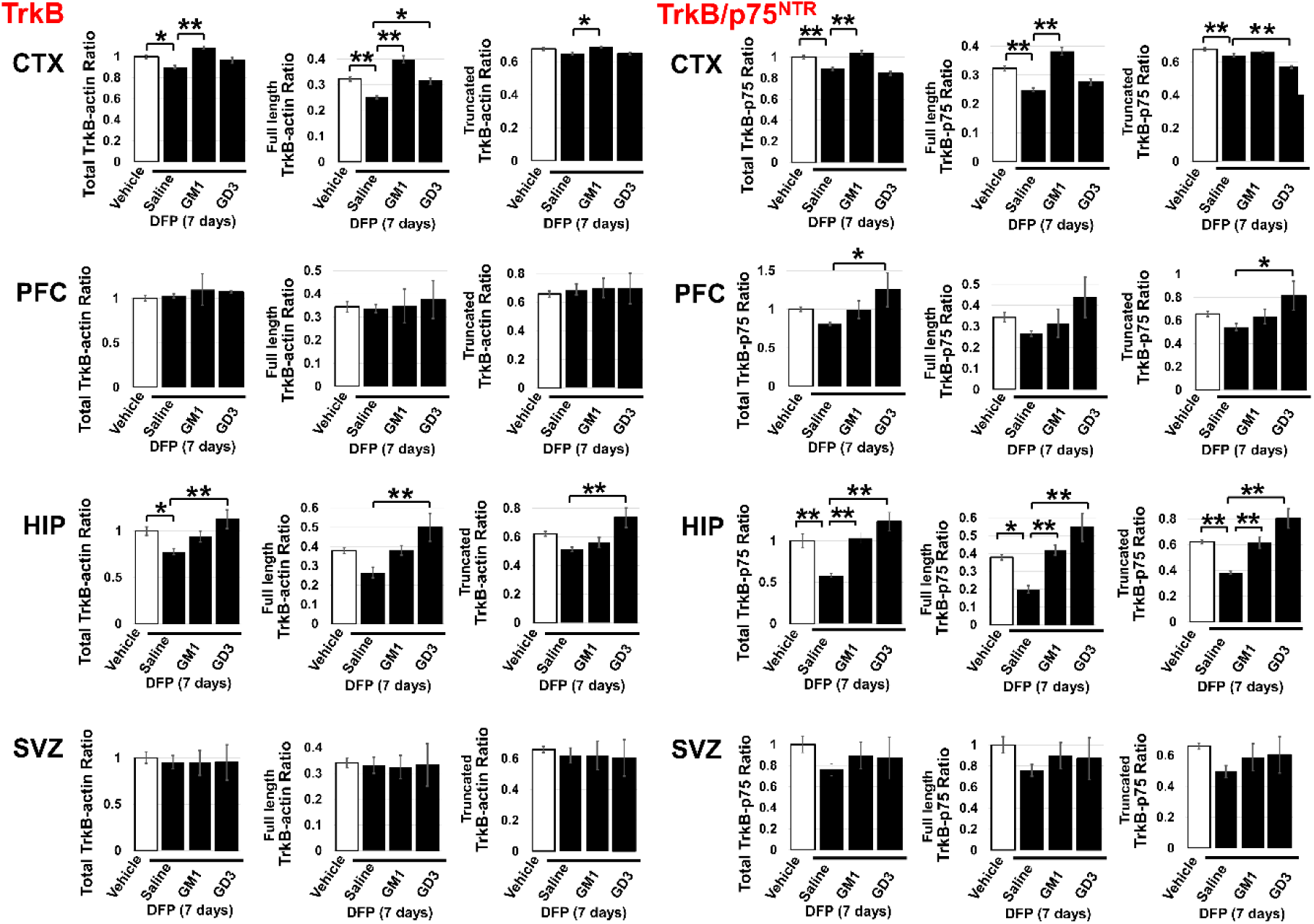
Quantification of Full-Length and Truncated TrkB. TrkB expression are reduced following DFP exposure (4.0 mg/kg) and restored by GM1 and GD3 treatments. Intranasally infused gangliosides (5 mg/kg/day for 7 days) ameliorate the decreased levels of TrkB in the cortex, prefrontal cortex, and hippocampus of the DFP-exposed mouse brain. Signals on immunoblots of Fig. 3A were quantified through image analysis. The columns represent the quantification of full-length, truncated, and total TrkB to actin (left) or p75^NT^(right) were calculated for each replicate sample. The TrkB/p75^NTR^ ratios were analyzed to determine whether a cell undergoes apoptosis or survives in the presence of neurotrophins. Results are means ± S.E.M. of ≥3 replicates per condition. *P < 0.05 and **P < 0.01 indicates significant differences between indicated groups, as calculated using one-way ANOVA with a Tukey’s multiple comparison test. CTX, cortex; PFC, prefrontal cortex; HIP, hippocampus; SVZ, subventricular zone.

**Fig. S4.**
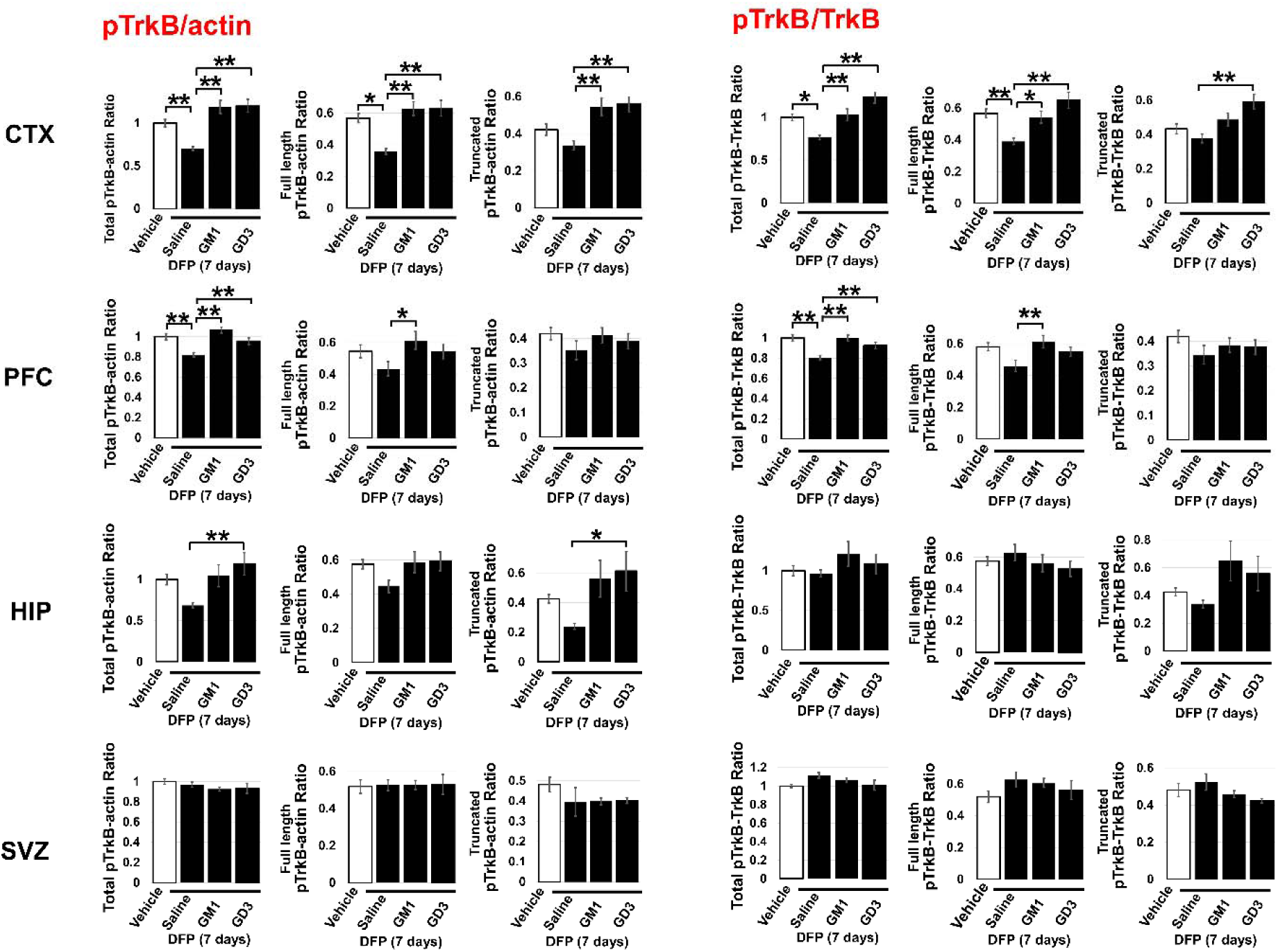
Quantification of Full-Length and Truncated TrkB Phosphorylation. TrkB phosphorylation are reduced following DFP exposure (4.0 mg/kg) and restored by GM1 and GD3 treatments. Intranasally infused gangliosides (5 mg/kg/day for 7 days) ameliorate the decreased posphorylation of TrkB in the cortex, prefrontal cortex, and hippocampus of the DFP-exposed mouse brain. Signals on immunoblots of Fig. 6A were quantified through image analysis. The columns represent the quantification of full-length, truncated, and total TrkB phosphorylation to actin (left; pTrkB/actin) or TrkB (right; pTrkB/TrkB) were calculated for each replicate sample. Results are means ± S.E.M. of ≥3 replicates per condition. *P < 0.05 and **P < 0.01 indicates significant differences between indicated groups, as calculated using one-way ANOVA with a Tukey’s multiple comparison test. CTX, cortex; PFC, prefrontal cortex; HIP, hippocampus; SVZ, subventricular zone.

**Fig. S5.**
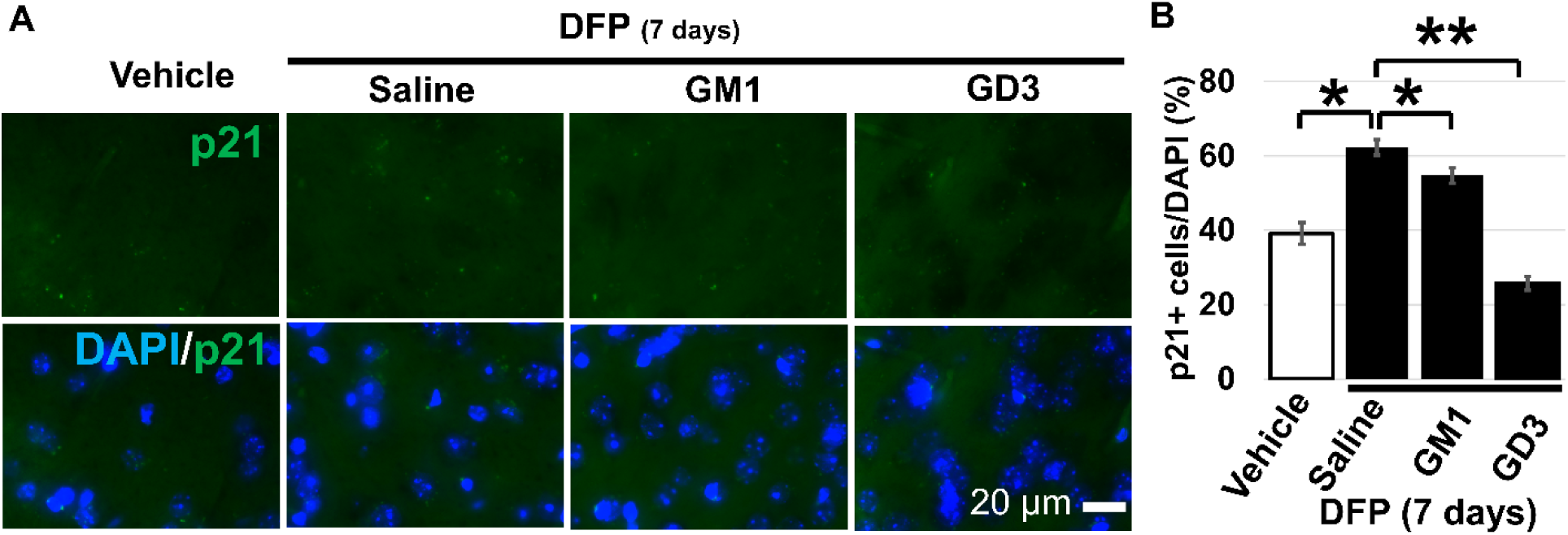
Restoration of the Senescence Marker p21 Levels in the Prefrontal Cortex of Mice Exposed to DFP through GM1 and GD3 Treatments. Intranasally infused gangliosides (5 mg/kg/day for 7 days) ameliorate the p21 expression after DFP exposure (4.0 mg/kg). Fluorescent images were acquired using a Cytation 3 cell imaging multi-mode microplate reader (Agilent Technologies) equipped with a 60x objective with identical acquisition settings. Gen5 software was used for initial image acquisition of fluorescent signals. (A) Fluorescent images for p21 (green) and the merged images with nuclear DAPI staining (blue). Scale bars = 20 μm. Bar plots illustrate the mean (± S.E.M.) percentage of cell expressing senescence marker p21 in prefrontal cortex. The results are based on quantifications from 3 replicates per condition.*P < 0.05 and **P < 0.01 indicates significant differences between indicated groups, as calculated using one-way ANOVA with a Tukey’s multiple comparison test.

